# BioML-bench: Evaluation of AI Agents for End-to-End Biomedical ML

**DOI:** 10.1101/2025.09.01.673319

**Authors:** Henry E. Miller, Matthew Greenig, Benjamin Tenmann, Bo Wang

**Author notes:** Contributed extensively to this work (Henry E. Miller, Matthew Greenig).

## Abstract

Large language model (LLM) agents hold promise for accelerating biomedical research and development (R&D). Several biomedical agents have recently been proposed, but their evaluation has largely been restricted to question answering (e.g., LAB-Bench) or narrow bioinformatics tasks. Presently, there remains a lack of benchmarks evaluating agent capability in multi-step data analysis workflows or in solving the machine learning (ML) challenges central to AI-driven therapeutics development, such as perturbation response modeling or drug toxicity prediction. We introduce **BioML-bench**, the first benchmarking suite for evaluating AI agents on end-to-end biomedical ML tasks. BioML-bench spans four domains (protein engineering, single-cell omics, biomedical imaging, and drug discovery) with tasks that require agents to parse a task description, build a pipeline, implement models, and submit predictions graded by established metrics (e.g., AUROC, Spearman). We evaluate four open-source agents: two biomedical specialists (STELLA, Biomni) and two generalists (AIDE, MLAgentBench). On average, agents underperform relative to human baselines, and biomedical specialization does not confer a consistent advantage. We also found that agents which employed more diverse ML strategies more often tended to score highest, suggesting that architecture and scaffolding may be stronger determinants of performance. These findings underscore both the potential and current limits of agentic systems for biomedical ML, and highlight the need for systematic, reproducible evaluations. BioML-bench is provided open-source at github.com/science-machine/biomlbench.

## 1 Introduction

Large language model (LLM) agents are systems built around LLMs that can plan and take actions autonomously via tools and code execution. They hold promise for transforming biomedical research and development (R&D) by automating time-consuming tasks such as literature search (Skarlinski et al. [2024]) and by enabling *in silico* experimentation, as exemplified by a recently-described virtual lab for identifying nanobodies against SARS-CoV-2 (Swanson et al. [2025]). A key opportunity is the reliable automation of biomedical machine learning (ML) workflows (e.g., perturbation-response modeling, image analysis, and drug property prediction) which demand competence across data preprocessing, model design/selection, and hyperparameter optimization. However, most evaluations of biomedical agents to date focus on question answering or narrow bioinformatics tasks (Jin et al. [2025], Huang et al. [2025]), leaving open the question of whether agents can complete multi-step, end-to-end ML pipelines that reflect real-world use cases.

We introduce **BioML-bench**, a benchmarking suite for evaluating AI agents on end-to-end biomedical ML tasks (Figure 1). BioML-bench comprises biomedical-specific tasks and metrics across four domains (protein engineering, single-cell omics, biomedical imaging, and drug discovery). To complete these tasks, agents must parse a task description, build data pipelines, implement models, and submit predictions, which are then graded by task-specific metrics (e.g., AUROC, Spearman). We evaluate two biomedical specialist agents (STELLA, Biomni) and two general-purpose ML agents (AIDE, MLAgentBench). While performance varies across agents, all agents on average underperform human baselines; moreover, we observe no consistent advantage for biomedical-specialized over generalist agents, suggesting that agent architecture and scaffolding may be the primary drivers of capability at present. In particular, we find that successful agents use more diverse ML strategies (e.g., feature engineering, model stacking) more often than less successful agents. Together, these findings clarify both the current reach and the limits of agentic systems for biomedical ML and motivate systematic, reproducible evaluation going forward. To meet this need, BioML-bench is released as a pip-installable package (with tutorials and a documentation website) in order to lower barriers for benchmarking and to accelerate reproducible progress in agentic biomedical ML: github.com/science-machine/biomlbench.

**Figure 1.**
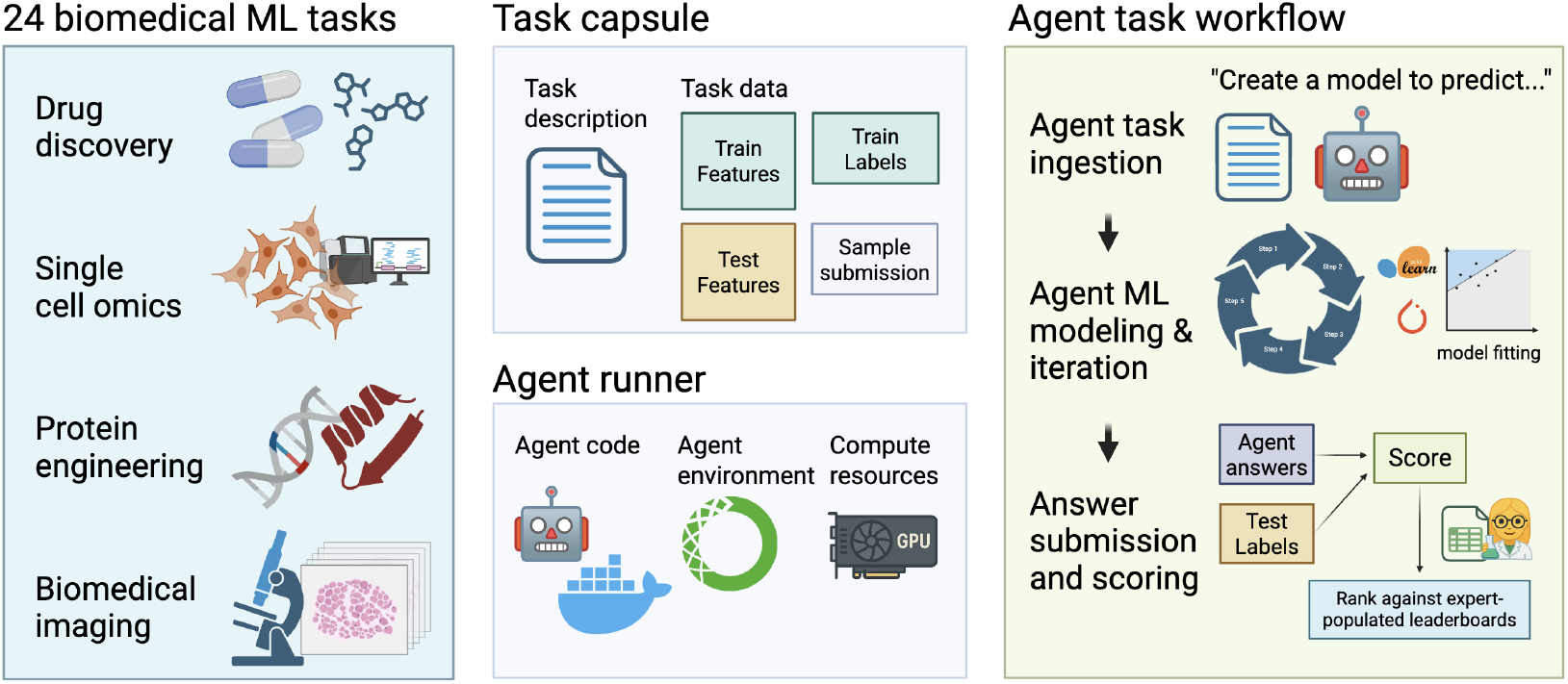
BioML-bench overview. The benchmark spans four biomedical ML domains (*protein engineering, single-cell omics, biomedical imaging, drug discovery*). Based upon MLE-bench, each task is packaged as a *task capsule* with a task description, training data, test features, and an example submission file. Agents run in a Docker container with packages provided via a pre-built conda environment. To complete a task, agents ingest the task description, design and fit ML models, and emit predictions that are automatically validated and graded by task-specific metrics (e.g., AUROC, Spearman) and ranked against human leaderboards which are largely expert-populated. Created with BioRender.com.

## 2 Related Work

### Benchmarks for Biomedical ML

Numerous biomedical benchmarks exist for evaluating ML models, including the Therapeutics Data Commons (drug discovery) (Huang et al. [2021]), ProteinGym (protein fitness prediction) (Notin et al. [2023]), BBBC (bioimage analysis) (Ljosa et al. [2012]), and OpenProblems (single-cell omics) (Luecken et al. [2025]). These provide valuable task definitions and human baselines, but they assess isolated model performance rather than whether autonomous agents can design and execute complete ML workflows.

### Biomedical Agents

Several biomedical agents have recently been proposed, including Biomni (Huang et al. [2025]), STELLA (Jin et al. [2025]), Virtual Lab (Swanson et al. [2025]), Perturbo-Agent (Hao et al. [2025]), CellForge (Tang et al. [2025]), and Cell Voyager (Alber et al. [2025]).

These systems highlight the potential of agentic AI for computational biology, spanning designs that emphasize self-improvement and orchestration of external tools. However, evaluations to date have typically been restricted to question answering, literature reasoning, or narrow domain-specific analyses such as perturbation panel design. For this work, we focused on STELLA and Biomni as the only biomedical agents sufficiently general to be evaluated across the variety of tasks in BioML-bench.

### Agent Benchmarks

Existing agent benchmarks are complementary, but major gaps remain. MLE-bench demonstrated that executable, end-to-end evaluation of ML agents is feasible at scale (Chan et al. [2025]). However, its scope is restricted to Kaggle-derived tasks, whose leaderboards are populated by crowd-sourced submissions rather than domain experts. BixBench provided a broad evaluation of bioinformatics agents with expert human baselines (Mitchener et al. [2025]), yet it relies mainly on text-based responses and bioinformatics tasks that do not involve supervised learning, leaving open whether agents can design and run complete biomedical ML workflows. Together, these efforts underscore the need for a benchmark that combines executable evaluation with biomedical domain-specific tasks and expert-populated leaderboards.

### Gap

To our knowledge, no existing benchmark evaluates whether LLM agents can complete end-to-end ML tasks across diverse biomedical domains. BioML-bench fills this gap by combining graded biomedical ML tasks with human baselines drawn mostly from expert-populated leaderboards. By reporting not only capability but also success and failure modes, it provides a more realistic measure of agent readiness for biomedical R&D.

## 3 Methods

### 3.1 Benchmark Setup

BioML-bench evaluates whether agents can design and execute full ML pipelines whose predictions are graded against a held-out ground truth. The framework is adapted from MLE-bench, but with key modifications to support biomedical contexts, including support for multiple biomedical task domains, new biomedical agents, tasks, and evaluation metrics, and support for bioinformatics-specific file formats (e.g., h5ad). It was also extended with multiple cloud deployment configurations and full mkdocs documentation. In BioML-bench, task capsules provide agents with a structured task description, training data, test features, and a sample submission in valid format. Agents must design, implement, and run ML models to generate test-set predictions, and emit outputs. Submissions are automatically validated and scored.

### 3.2 Task Suite

The benchmark spans four biomedical ML domains central to R&D: drug discovery, biomedical imaging, protein engineering, and single-cell omics. Tasks were selected for both practical relevance to biotechnology and the availability of populated human leaderboards to provide performance baselines. Sources include PolarisHub (drug discovery), Kaggle (biomedical imaging), ProteinGym (protein engineering), and OpenProblems (single-cell) (Table 1, with detailed task descriptions in Table 3).

**Table 1:** Summary of task domains and sources. Representative tasks are shown; full task lists appear in the appendix.

| Domain | Representative tasks | Source |
| --- | --- | --- |
| Drug discovery | ADME/Tox property prediction (solubility, BBB, CYP2D6) | PolarisHub |
| Biomedical imaging | Cancer detection, tumor segmentation, fibrosis progression | Kaggle |
| Protein engineering | Activity prediction, stability prediction | ProteinGym |
| Single-cell omics | Multimodal prediction, perturbation response, cell-cell communication | OpenProblems |

### 3.3 Agents

We evaluated four agent frameworks: two biomedical research agents (Biomni (Huang et al. [2025]) and STELLA (Jin et al. [2025])) and two general-purpose ML agents (AIDE (Jiang et al. [2025]) and MLAgentBench (Huang et al. [2024])). Biomni (specifically, v0.0.5) and STELLA were chosen as the only biomedical agents sufficiently general to operate across multiple task domains, while AIDE (v6.3.3) and MLAgentBench represent strong baselines from MLE-bench. At evaluation time, we used the latest compatible LLM backends: GPT-4.1 for MLAgentBench and AIDE, Claude Sonnet 4 for Biomni, and the OpenRouter default for STELLA. A simple “Dummy” agent was also included, producing uninformative predictions (random or zero depending on the task).

### 3.4 Evaluation Protocol and Compute Budget

Each task provides a structured task card describing the objective and dataset. Agents must load data, build pipelines, train models, and output predictions to a file in a standardised format. Submissions are validated and graded using the task’s original metric. For each agent–task combination, four independent replicates were run. To quantify both performance and reliability we report:

- **Leaderboard percentile:** agent score mapped to the public leaderboard percentile for the task (*higher is better*). Runs with no valid submission either receive a percentile of zero (penalized scoring) or are not considered (non-penalized scoring).
- **Mean rank:** average relative rank of an agent (compared to other agents) across tasks, based on leaderboard percentile (*lower is better*). This allows aggregation across domains with varying leaderboard distributions.
- **Above-median (%):** proportion of tasks where the agent’s mean score exceeds the median human submission.
- **Any-medal (%):** proportion of tasks where the agent achieves a public leaderboard medal threshold, following the MLE-bench definition (Chan et al. [2025]).
- **Completion rate (%):** proportion of tasks that yield a valid, gradable submission.

All runs were executed under matched resource budgets to ensure comparability. Imaging tasks used a single NVIDIA L4 GPU (16 vCPUs, 64 GB RAM) with a 16-hour wall-clock limit. All other tasks used CPU-only machines with identical specifications and an 8-hour limit.

## 4 Results

### 4.1 Overall performance

Table 2 reports aggregate agent performance across all tasks. In the “penalized” setting, failed runs (runs in which an agent fails to produce a valid submission file) are assigned a leaderboard percentile of zero, reflecting the view that a failed run corresponds to the poorest possible performance. Under this analysis, Biomni (which had no failed tasks) achieves the strongest overall results, with the best leaderboard percentile (34.45±5.36),= and mean rank (1.88 ±0.18). In the “non-penalized” setting, failed runs are excluded from percentile calculations, reflecting the idea that many failures stem from scaffolding issues rather than fundamental capability and thus shouldn’t penalize benchmarking performance. Here, STELLA is highly competitive with Biomni, outperforming it on most metrics.

**Table 2:** Overall BioML-bench performance. Metrics defined in Section 3.4. In the penalized setting, failed runs are scored as zero; in the non-penalized setting, failed runs are omitted. Biomni achieves the highest penalized performance while STELLA achieves the highest in the non-penalized regime. However, the performance difference between AIDE, STELLA, and Biomni is generally within the reported SEM.

| Type | Agent | Leaderboard Percentile | Mean Rank | Above Median (%) | Any Medal (%) | Completion Rate (%) |
| --- | --- | --- | --- | --- | --- | --- |
| Penalized | Dummy | 2.50 ± 1.16 | 4.33 ± 0.21 | 0.0 ± 0.0 | 0.0 ± 0.0 | **100.0** |
| | MLAgentBench | 21.43 ± 4.40 | 3.06 ± 0.24 | 16.7 ± 5.6 | 6.2 ± 3.4 | 97.9 |
| | AIDE | 28.50 ± 5.22 | 2.56 ± 0.23 | 28.1 ± 6.4 | 12.5 ± 5.2 | 80.2 |
| | STELLA | 31.84 ± 5.82 | 2.62 ± 0.24 | **34.4 ± 7.6** | 17.7 ± 6.1 | 91.7 |
| | Biomni | **34.45 ± 5.36** | **1.88 ± 0.18** | 32.3 ± 8.0 | **19.8 ± 6.7** | **100.0** |
| Non-penalized | Dummy | 2.50 ± 1.16 | 4.38 ± 0.20 | 0.0 ± 0.0 | 0.0 ± 0.0 | - |
| | MLAgentBench | 23.44 ± 4.65 | 3.07 ± 0.19 | 18.5 ± 5.9 | 7.6 ± 4.0 | - |
| | AIDE | 37.41 ± 5.85 | 2.20 ± 0.21 | 36.2 ± 7.9 | 14.5 ± 6.0 | - |
| | STELLA | **39.45 ± 6.52** | 2.45 ± 0.21 | **42.8 ± 8.7** | **20.8 ± 7.1** | - |
| | Biomni | 34.45 ± 5.36 | **2.08 ± 0.22** | 32.3 ± 8.0 | 19.8 ± 6.7 | - |

### 4.2 Domain-level analysis of performance

Figure 2 and Table 4 summarize agent performance by domain with penalties applied. In single-cell omics, Biomni is strongest, closely followed by STELLA and AIDE. In Biomedical Imaging, MLAgentBench yields the best mean percentile, while in Protein Engineering and Drug Discovery Biomni leads. Removing the failed-run penalty changes the picture: AIDE and STELLA appear more robust, with rankings shifting across domains. Specifically, STELLA leads Biomedical Imaging by a wide margin, AIDE leads Protein Engineering, and Biomni leads Drug Discovery. As with overall results, domain-level analyses indicate that once failure penalties are removed, Biomni no longer shows a consistent performance lead compared to the other agents.

**Figure 2.**
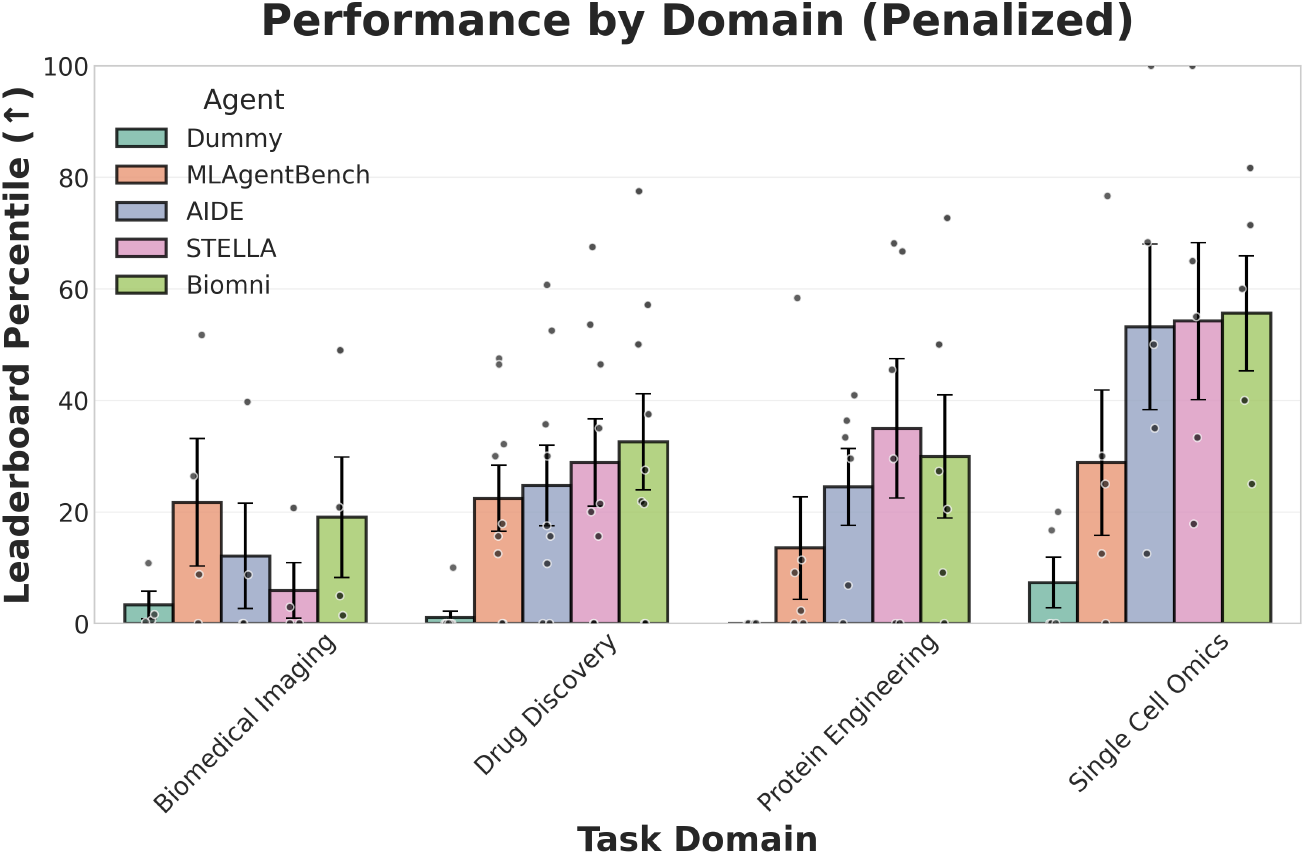
Domain-level agent performance (penalized). Per-task leaderboard percentiles by agent within each domain. Points are individual tasks; bars and whiskers summarize the mean±SEM. Failed run penalties are applied such that a run without a successful answer submission is graded as having a leaderboard percentile of zero.

### 4.3 Successful Agent Strategies

To better understand factors driving performance, we manually examined logs from runs where agents exceeded the human leaderboard median (percentile >50%). Across 94 such runs, we cataloged the prevalence of different ML strategies (Figure 3). The strongest agents overall (AIDE, STELLA,

**Figure 3.**
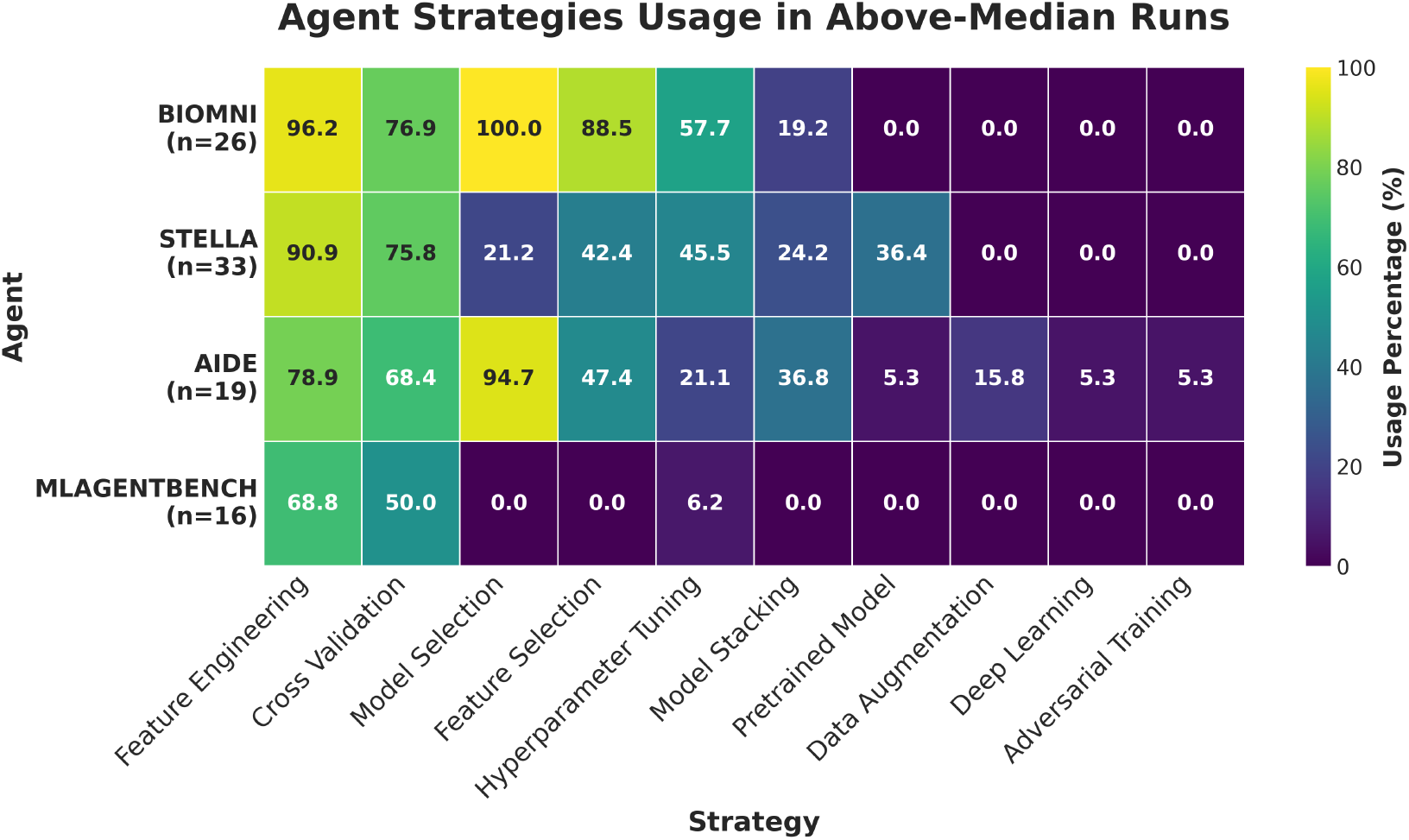
ML strategies used by agents in above-median runs. Heatmap showing the proportion of above-median runs (runs in which agents perform above 50% of the human leaderboard) in which an agent used any particular ML strategy.

and Biomni) employed a wider variety of strategies more often, frequently testing multiple feature engineering approaches and model types. Detailed examination of particularly successful runs sometimes revealed sophisticated agent strategies leveraging clear domain understanding, even for generalist agents. For example, AIDE achieved a 100th percentile leaderboard position on the Tsuboyama 2023 ProteinGym benchmark (Notin et al. [2023], Tsuboyama et al. [2023]) using advanced feature engineering approaches that involved calculating a wide range of evolutionary and biophysical quantities from raw protein sequences, including BLOSUM substitution scores, sequence conservation indices, and specific biochemical characteristics of amino acids such as charge, volume, and hydrogen bond propensities. Interestingly, these strategies were not always consistently applied, even in replicate runs with the same agent on the same task; in fact, on a different run for the Tsuboyama task, AIDE achieved only a 54th percentile leaderboard position using an approach with a significantly less-extensive protein feature engineering pipeline.

Substantial variability was also observed in the ML architectures chosen by agents (Figure 5): Biomni and MLAgentBench relied on random forests in roughly 50% of cases; STELLA alternated among random forests, ridge regression, and XGBoost; AIDE often used stacked or gradient boosting models (e.g., LightGBM). Strikingly, deep learning (DL) was rarely attempted and almost never selected as the final model, even on image-based tasks. This stands in contrast to human leaderboards, which are dominated by DL approaches.

### 4.4 Agent failure modes

We also analyzed 41 failed runs to characterize agent failure modes (Figure 6). AIDE often failed due to resource exhaustion, particularly when copying large image files. MLAgentBench and STELLA frequently exited early without producing outputs, often due to scripts that executed but failed to save predictions (e.g., silent errors from try–except fallbacks) or agents attempting unnecessary environment creation that stalled on package installation. AIDE and STELLA were affected by uncaught exceptions in their frameworks. Finally, some runs exceeded the time limit without producing results. Importantly, these failure modes did not reflect fundamental architectural flaws but rather are likely addressable issues in scaffolding and execution.

## 5 Discussion

Our results show that both specialized biomedical agents and general-purpose ML agents can solve end-to-end tasks in biology-heavy domains such as single-cell omics and protein engineering. While Biomni achieved the strongest average performance, STELLA overtook it once failed-run penalties were removed, suggesting that scaffolding issues (e.g., resource exhaustion, uncaught exceptions) rather than fundamental agent limitations are an important factor in benchmarking performance. Moreover, while Biomni and STELLA demonstrated strong benchmarking performance, their advantage over one non-specialist agent, AIDE, was typically within the SEM range. These findings indicate that biomedical specialization alone does not guarantee superior performance on biomedical ML tasks. More broadly, all agents struggled to consistently exceed median human leaderboard performance, with Biomni, STELLA, and AIDE averaging only in the 34–39th percentile range.

Analysis of successful runs revealed that top-performing agents primarily relied on classical ML strategies such as feature engineering, feature selection, and hyperparameter tuning. Strikingly, these approaches achieved strong leaderboard placements despite the fact that human submissions were often dominated by deep learning models. Failure analysis was equally revealing: most failed runs were attributable to scaffolding issues (e.g., silent errors in agent-written code or uncaught framework exceptions) rather than inherent weaknesses of the underlying agent architectures. Taken together, these results underscore both the reach and the current limits of agentic systems for biomedical ML. Agents are capable of assembling and executing workable pipelines across diverse biomedical tasks, but they generally underperform human domain experts, and their reliability is heavily constrained by framework issues.

While BioML-bench reuses the scaffold of MLE-bench (e.g., task capsules, automated grading), it diverges in scope and substance: tasks are drawn from ProteinGym, OpenProblems, and PolarisHub rather than just from Kaggle, extended to biomedical data formats, and benchmarked against expert-populated leaderboards. These differences shift the evaluation from ranking well in generic ML competitions to solving scientific ML problems end-to-end, better-reflecting the task domains and constraints of real-world biomedical R&D.

Overall, BioML-bench provides the community with a reproducible and extensible framework for evaluating agentic systems on real biomedical ML problems. By grounding evaluation in verifiable outputs rather than textual answers, it enables systematic progress toward agents that can automate meaningful components of computational biomedical research. Future work will expand the task suite, incorporate additional agents, and investigate strategies for improving both capability and robustness, with the long-term goal of advancing trustworthy agentic systems for biomedical R&D.

### 5.1 Limitations and Future Work

Some limitations of our study should be noted. First, the choice of LLMs for agents introduces potential confounds: stronger base LLMs may yield stronger agent performance. To mitigate this, each agent was paired with its recommended or default LLM (using the latest-generation compatible version). Still, future work should systematically vary LLMs to more cleanly disentangle agent design from LLM capability.

Additionally, the interpretation of human performance is itself imperfect. Public leaderboards inevitably reflect a mix of participant expertise, effort, and compute budgets. A key advantage of the biomedical benchmarks used in BioML-bench is that their leaderboards are populated largely by domain experts, providing a more meaningful point of reference than generic crowd-sourced competitions. Still, our use of these leaderboards remains pragmatic: they offer an external, reproducible baseline for situating agent performance, consistent with MLE-bench. Future work may consider complementary baselines, such as automated ML optimization systems like H2O AutoML (LeDell and Poirier [2020]) and Lazy Predict, which are useful for evaluating whether LLM agents that employ classical ML approaches provide an advantage over automated pipelines. It will also be valuable to explore agent scaffolding that better supports deep learning architectures, as current agents did not seem willing to use these approaches. Finally, it will be interesting to evaluate the impact of tool-usage (e.g., web search, literature review), memory/self-evolving agents, and multi-agent system designs on agent performance.

## Acknowledgments

Gabriel Mejia provided extensive and valuable feedback on this manuscript. Additionally, ScienceMachine provided the compute and LLM API resources for this work.

## A Supplementary Material

**Table 3:**
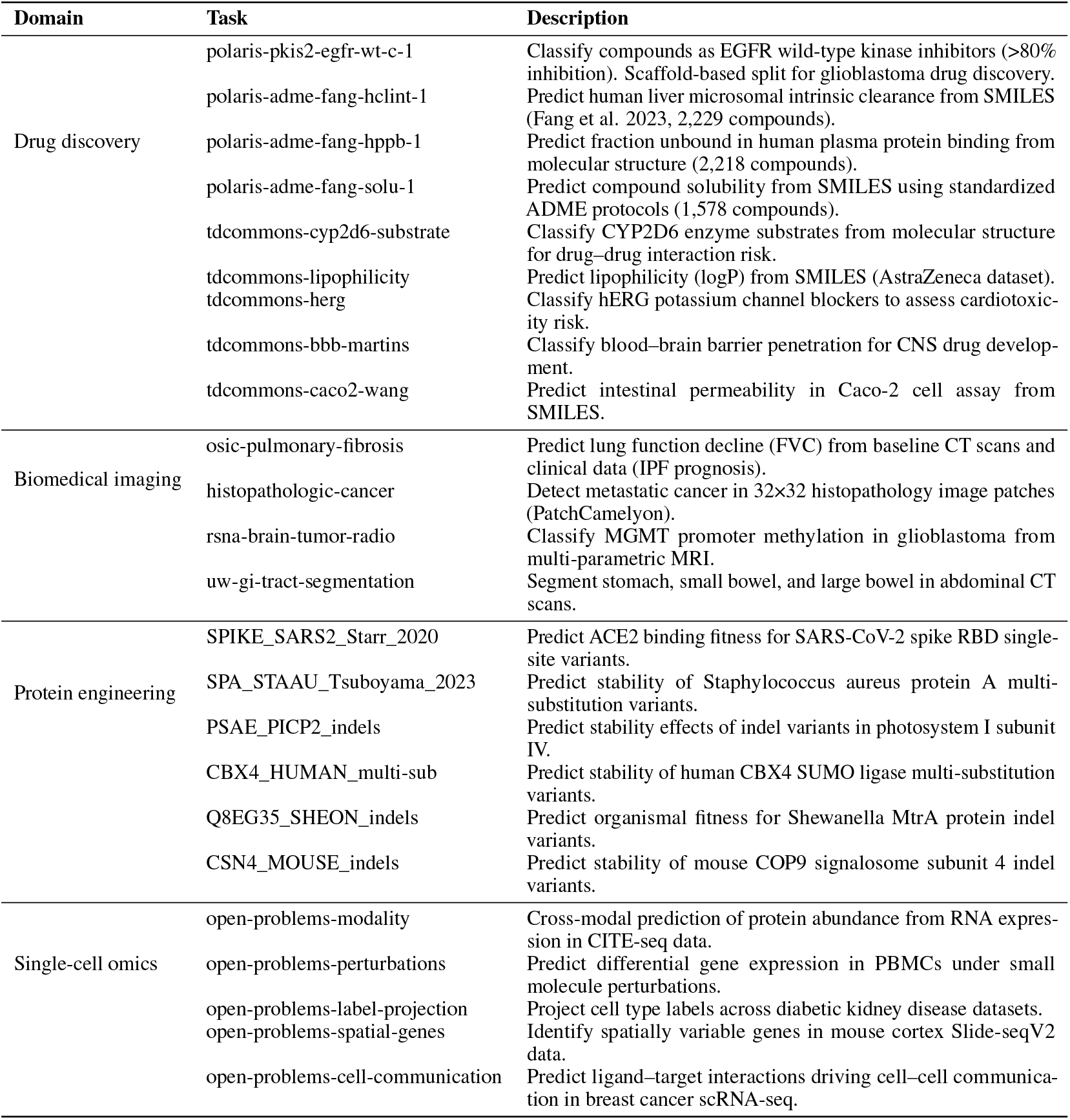
Comprehensive task descriptions for the BioML-bench suite. Each row specifies the domain, task identifier, and biomedical objective, with dataset sources noted in the text. This table provides the full task set underlying BioML-bench; a high-level summary of domains and representative problems is given in Table 1.

**Figure 4.**
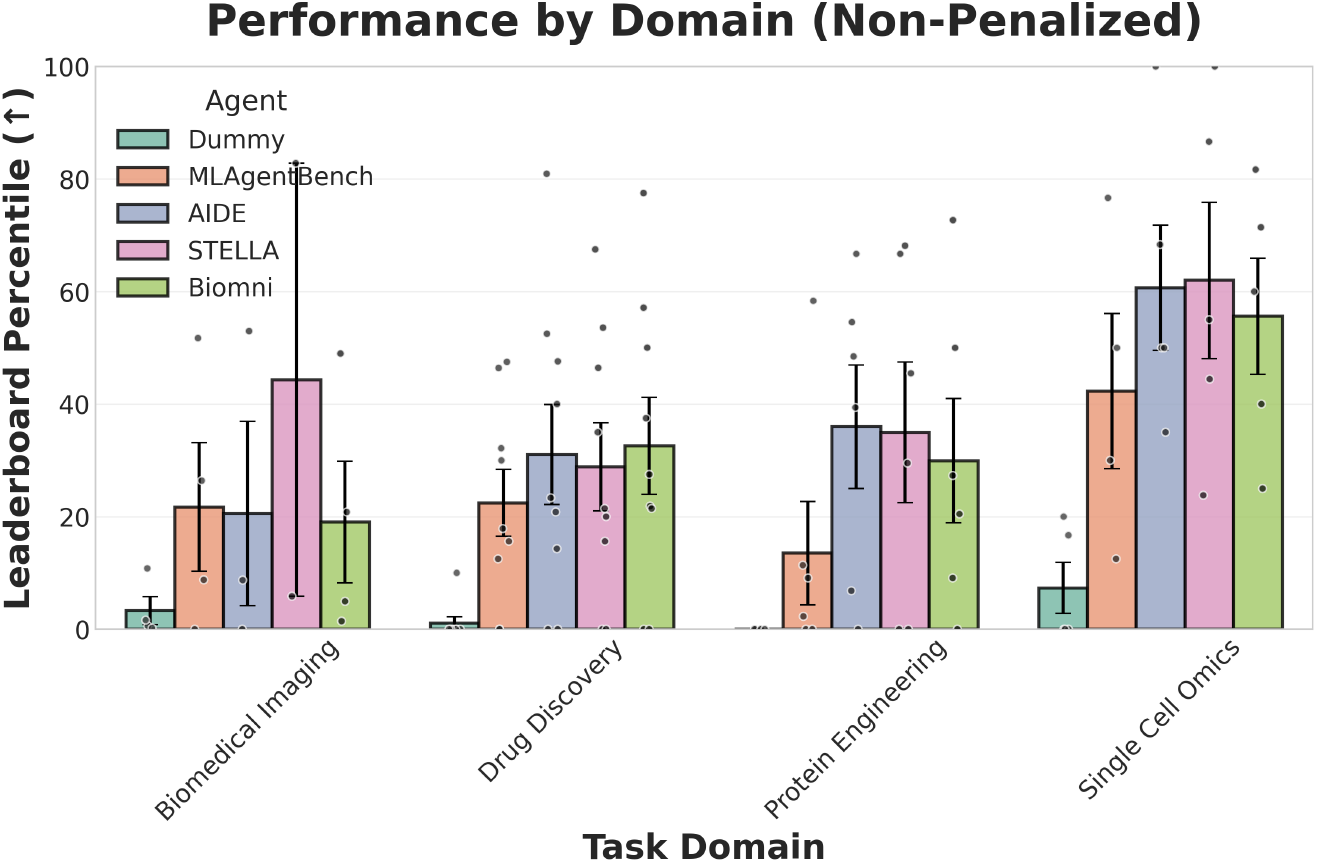
Domain-level agent performance in the non-penalized setting. Each point shows the leaderboard percentile for a single task; bars and whiskers indicate mean ± SEM across tasks within a domain. Unlike Figure 2, failed runs are omitted rather than assigned a score of zero.

**Figure 5.**
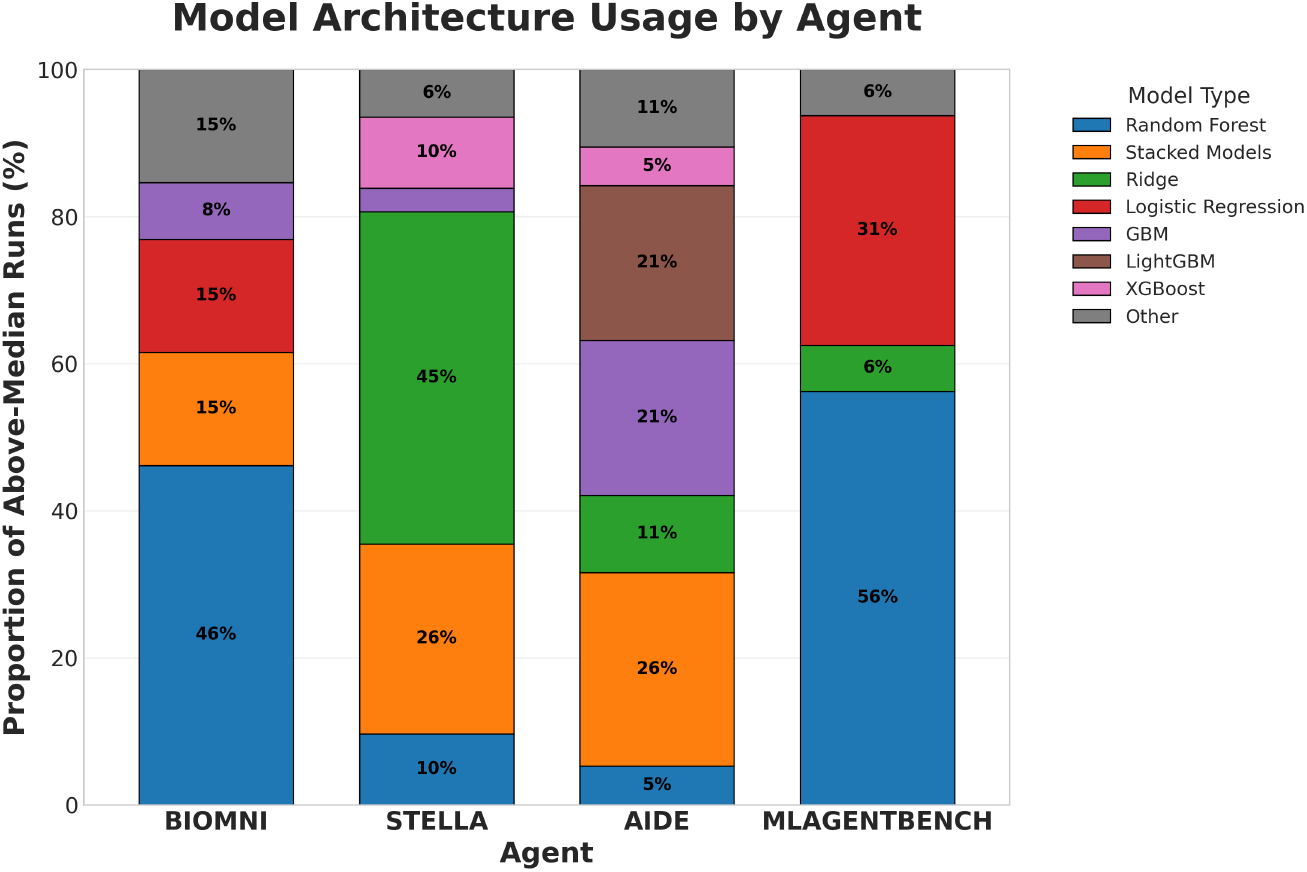
Model architectures used by agents in above-median runs (i.e., runs scoring above 50% of the human leaderboard). Bars show the proportion of runs using each architecture. “Stacked Models” denotes any metalearner that combines multiple base models.

**Table 4:** BioML-bench results by domain. Metrics are defined in Section 3.4. Results are reported under penalized (failed runs scored as zero) and non-penalized (failed runs omitted) settings.

| Type | Domain | Agent | Leaderboard Percentile | Mean Rank | Above Median (%) | Any Medal (%) | Completion Rate (%) |
| --- | --- | --- | --- | --- | --- | --- | --- |
| Penalized | Biomedical Imaging | Dummy | 3.30 ± 2.52 | 3.50 ± 0.87 | 0.0 ± 0.0 | 0.0 ± 0.0 | NA |
| | | MLAgentBench | **21.73 ± 11.41** | **2.00 ± 0.71** | **12.5 ± 12.5** | 0.0 ± 0.0 | **100.0** |
| | | AIDE | 12.11 ± 9.43 | 3.00 ± 0.91 | 6.2 ± 6.2 | 0.0 ± 0.0 | 81.2 |
| | | STELLA | 5.91 ± 4.98 | 4.00 ± 0.41 | 6.2 ± 6.2 | 0.0 ± 0.0 | 68.8 |
| | | Biomni | 19.04 ± 10.83 | 2.50 ± 0.29 | **12.5 ± 12.5** | **12.5 ± 12.5** | **100.0** |
| | Drug Discovery | Dummy | 1.11 ± 1.11 | 4.44 ± 0.28 | 0.0 ± 0.0 | 0.0 ± 0.0 | NA |
| | | MLAgentBench | 22.45 ± 5.93 | 2.83 ± 0.20 | 16.7 ± 9.3 | 5.6 ± 3.7 | **100.0** |
| | | AIDE | 24.75 ± 7.23 | 2.50 ± 0.33 | 25.0 ± 11.0 | 5.6 ± 5.6 | 80.6 |
| | | STELLA | 28.84 ± 7.84 | 2.39 ± 0.26 | 25.0 ± 11.0 | **13.9 ± 7.3** | **100.0** |
| | | Biomni | **32.55 ± 8.61** | **1.67 ± 0.29** | **30.6 ± 12.3** | 13.9 ± 11.1 | **100.0** |
| | Protein Engineering | Dummy | 0.00 ± 0.00 | 4.50 ± 0.26 | 0.0 ± 0.0 | 0.0 ± 0.0 | NA |
| | | MLAgentBench | 13.52 ± 9.18 | 3.50 ± 0.52 | 12.5 ± 12.5 | 0.0 ± 0.0 | **100.0** |
| | | AIDE | 24.50 ± 6.90 | 2.25 ± 0.44 | 25.0 ± 9.1 | 12.5 ± 8.5 | 75.0 |
| | | STELLA | **34.98 ± 12.51** | 2.33 ± 0.57 | **45.8 ± 18.7** | 16.7 ± 12.4 | **100.0** |
| | | Biomni | 29.93 ± 11.04 | **2.00 ± 0.45** | 25.0 ± 17.1 | **25.0 ± 17.1** | **100.0** |
| | Single Cell Omics | Dummy | 7.34 ± 4.52 | 4.60 ± 0.40 | 0.0 ± 0.0 | 0.0 ± 0.0 | NA |
| | | MLAgentBench | 28.83 ± 13.04 | 3.80 ± 0.49 | 25.0 ± 13.7 | 20.0 ± 14.6 | 90.0 |
| | | AIDE | 53.17 ± 14.86 | 2.70 ± 0.44 | 55.0 ± 16.6 | 35.0 ± 18.7 | 85.0 |
| | | STELLA | 54.23 ± 14.09 | 2.30 ± 0.54 | **60.0 ± 15.0** | **40.0 ± 20.3** | 85.0 |
| | | Biomni | **55.62 ± 10.32** | **1.60 ± 0.40** | **60.0 ± 20.3** | 30.0 ± 14.6 | **100.0** |
| Non-penalized | Biomedical Imaging | Dummy | 3.30 ± 2.52 | 3.50 ± 0.87 | 0.0 ± 0.0 | 0.0 ± 0.0 | - |
| | | MLAgentBench | 21.73 ± 11.41 | 2.12 ± 0.52 | 12.5 ± 12.5 | 0.0 ± 0.0 | - |
| | | AIDE | 20.56 ± 16.40 | 2.17 ± 0.73 | 11.1 ± 11.1 | 0.0 ± 0.0 | - |
| | | STELLA | **44.32 ± 38.47** | **2.00 ± 1.00** | **50.0 ± 50.0** | 0.0 ± 0.0 | - |
| | | Biomni | 19.04 ± 10.83 | 3.00 ± 0.41 | 12.5 ± 12.5 | **12.5 ± 12.5** | - |
| | Drug Discovery | Dummy | 1.11 ± 1.11 | 4.44 ± 0.28 | 0.0 ± 0.0 | 0.0 ± 0.0 | - |
| | | MLAgentBench | 22.45 ± 5.93 | 2.89 ± 0.20 | 16.7 ± 9.3 | 5.6 ± 3.7 | - |
| | | AIDE | 31.06 ± 8.88 | 2.33 ± 0.29 | **30.6 ± 13.3** | 7.4 ± 7.4 | - |
| | | STELLA | 28.84 ± 7.84 | 2.78 ± 0.26 | 25.0 ± 11.0 | **13.9 ± 7.3** | - |
|  |  | Biomni | **32.55 ± 8.61** | **1.67 ± 0.29** | **30.6 ± 12.3** | **13.9 ± 11.1** | - |
| | Protein Engineering | Dummy | 0.00 ± 0.00 | 4.50 ± 0.26 | 0.0 ± 0.0 | 0.0 ± 0.0 | - |
| | | MLAgentBench | 13.52 ± 9.18 | 3.67 ± 0.36 | 12.5 ± 12.5 | 0.0 ± 0.0 | - |
| | | AIDE | **35.99 ± 10.96** | **1.92 ± 0.45** | 38.9 ± 15.9 | 16.7 ± 11.4 | - |
| | | STELLA | 34.98 ± 12.51 | 2.58 ± 0.54 | **45.8 ± 18.7** | 16.7 ± 12.4 | - |
| | | Biomni | 29.93 ± 11.04 | 2.00 ± 0.45 | 25.0 ± 17.1 | **25.0 ± 17.1** | - |
| | Single Cell Omics | Dummy | 7.34 ± 4.52 | 4.80 ± 0.20 | 0.0 ± 0.0 | 0.0 ± 0.0 | - |
| | | MLAgentBench | 42.29 ± 13.78 | 3.50 ± 0.29 | 37.5 ± 16.1 | 31.2 ± 18.8 | - |
| | | AIDE | 60.67 ± 11.16 | 2.30 ± 0.54 | 58.3 ± 16.7 | 33.3 ± 19.0 | - |
|  |  | STELLA | **61.97 ± 13.91** | **1.90 ± 0.33** | **68.3 ± 15.0** | **46.7 ± 22.6** | - |
| | | Biomni | 55.62 ± 10.32 | 2.20 ± 0.58 | 60.0 ± 20.3 | 30.0 ± 14.6 | - |

**Figure 6.**
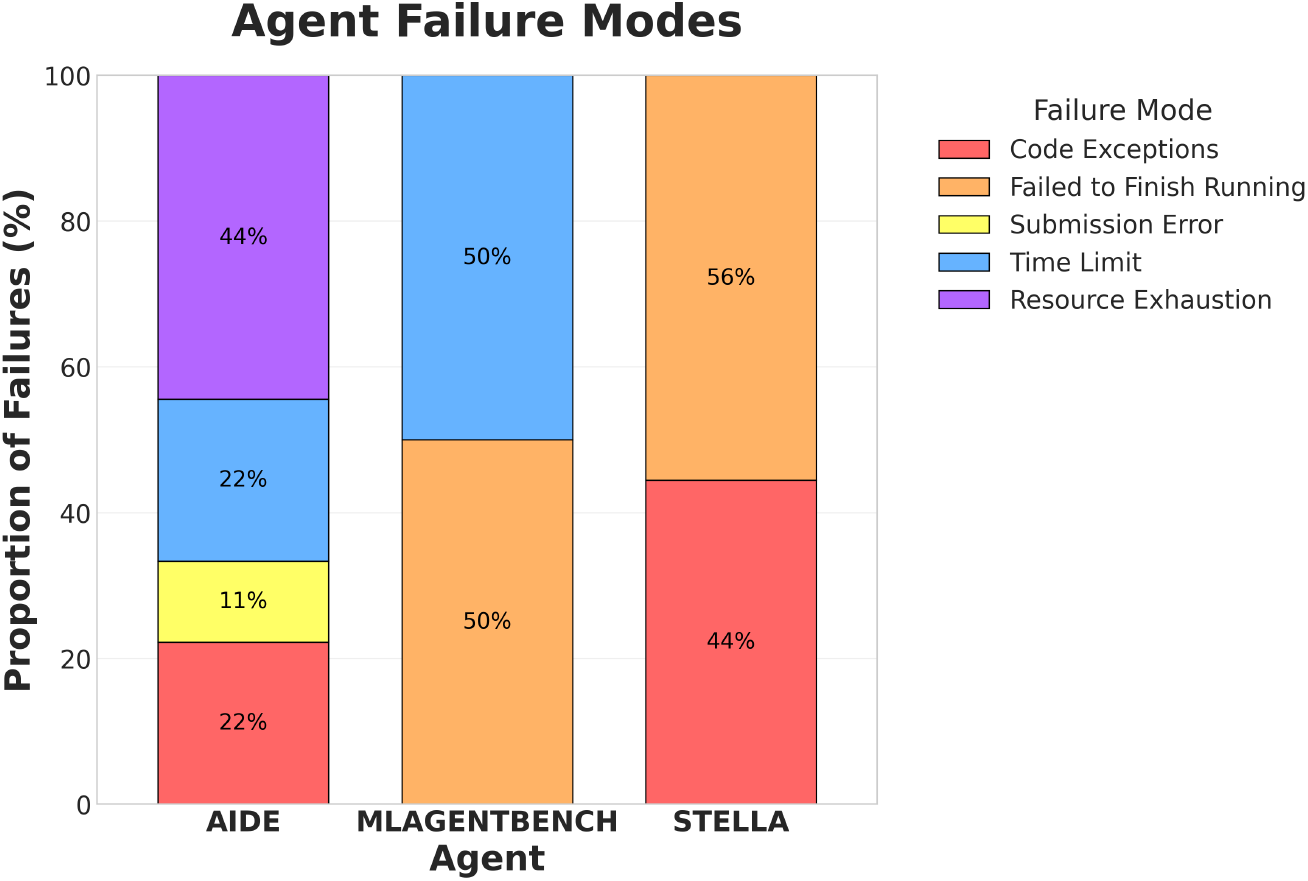
Failure mode analysis. Stacked bar chart showing the proportion of failed runs for each agent attributable to specific failure modes.

